# A network of mixed actin polarity in the leading edge of spreading cells

**DOI:** 10.1101/2022.08.26.505326

**Authors:** Wen-Lu Chung, Matthias Eibauer, Wenhong Li, Rajaa Boujemaa-Paterski, Benjamin Geiger, Ohad Medalia

**Affiliations:** Department of Biochemistry, University of Zurich, Winterthurerstrasse 190, 8057 Zurich, Switzerland; Department of Immunology, and regenerative Biology, Weizmann Institute of Science, Rehovot 76100, Israel

## Abstract

Physical interactions of cells with the underlying extracellular matrix (ECM) play key roles in multiple cellular processes, such as tissue morphogenesis, cell motility, wound healing, cancer invasion and metastasis. The actin cytoskeletal network is a central driver and regulator of cellular dynamics, that produces membrane protrusions such as lamellipodia and filopodia. In this study, we examined actin organization in the expanding lamellipodia during the early stages of cell spreading. To gain insight into the 3D actin organization, at a molecular resolution, we plated cultured fibroblasts on galectin-8 coated EM grids, an ECM protein presents in disease states. We then combined cryo-electron tomography (cryo-ET) with advanced image processing tools for reconstructing the structure of F-actin in the lamellipodia. This approach enabled us to resolve the polarity and orientation of the filaments, and the structure of the Arp2/3 complexes associated with F-actin branches. We show here that F-actin in lamellipodial protrusions forms a dense network with three distinct sub-domains. One consists primarily of radial filaments, with their barbed ends pointing towards the membrane, the other is enriched with parallel filaments that run between the radial fibers, in addition to an intermediate sub-domain. Surprisingly, a minor, yet significant (∼10%) population of actin filaments, are oriented with their barbed ends towards the cell center. Our results provide novel structural insights into F-actin assembly and dynamic reorganization in the leading edge of spreading cells.

## Introduction

Cell interactions with the extracellular matrix (ECM) play key roles in multiple cellular processes including tissue coherence and morphogenesis, cell migration, cell survival and cytoskeletal organization ^1-4^. Adhesive interactions with the matrix, primarily those mediated by integrin receptors, induce a local assembly of the actin cytoskeleton, that produces and coordinates both protrusive and contractile responses that drive cell motility ^5^. Initial contacts between cells and the underlying ECM lead to progressive cell spreading, driven by major deformations of the membrane ^6^. At the molecular level, the early integrin-mediated adhesive interactions initiate a complex cascade of cytoskeletal assembly and signaling events. Concertedly, they trigger radial cell spreading ^7, 8^, that is usually followed by cell polarization, manifested by the generation of a protrusive leading edge and a contractile trailing edge, at the interior and posterior aspects of the cell, respectively ^9-11^.

Actin filaments polymerize at the leading edge and apply mechanical forces that drive the protrusive extension of the plasma membrane ^12, 13^. Together with diverse actin-associated proteins, they form a robust skeletal network that displays a retrograde flow, regulated by a fine balance between actin polymerization at the front and interaction with nascent matrix adhesions, at the lamellipodium-lamella junction ^14-16^. Importantly, the Arp2/3 complex plays a key role in the formation and mechanics of the lamellipodium, by nucleating actin polymerization and branching ^17-19^. The lamellipodium of adherent cells is a sheet-like protrusion that is typically hundreds of nanometers in thickness ^20^. In migratory fibroblasts, the lamellipodium undergoes cycles of polymerization-dependent protrusion and acto-myosin driven retraction, that are believed to be regulated by variations in the mechanical load generated by the F-actin network ^21-23^.

The organization of actin filaments within the lamellipodium was extensively studied by light and electron microscopy. These studies confirmed the presence of a branched F-actin network and suggested that the vast majority of actin filaments at the leading edge direct their barbed ends toward the plasma membrane ^24-26^. However, these studies were mostly based on the use of permeabilized or fixed cell models that provided limited information concerning the precise filament organization and polarity.

The mode of cell spreading on the ECM is profoundly affected by the molecular composition of the underlying matrix and the set of adhesion receptors the cells possess ^27, 28^. Commonly, cell adhesion and spreading studies were conducted using specific adhesive proteins such as fibronectin, vitronectin, collagen and laminin ^29-32^. Each of these proteins may present different chemical and physical properties, interact with a different set of membrane receptors, and thus display distinct effects on cell spreading and motility. Distinct adhesive features were reported for members of the galectin family ^33^, comprising of several adhesive galactoside-binding animal lectins, that interact with a variety of cell-surface glycans. They are expressed in a variety of tissues and were shown to affect cell-ECM and cell-cell adhesion, and affecting trans-membrane signaling, cell spreading and cell migration ^34^. Most importantly, galectins contribute to cancer progression by regulating the migration and cell adhesion properties of tumor cells. ^35^

Within the galectin family, the dimeric galectin-8 (gal-8) was shown to play an important role in platelets activation and angiogenesis ^36^ and induce fast and efficient spreading of cells ^37^. Moreover, it was recently demonstrated that cell adhesion to gal-8-coated surfaces leads to essentially continuous spreading dynamics, unlike spreading on fibronectin, which consists of protrusion-and-retraction cycles ^38^. Consequently, the average projected area of fully spread HeLa cells, on gal-8 is about twice larger than the projected area of the same cells plated on fibronectin ^16^. It is notable that the initial spreading of the cells on gal-8 is radial, forming thin lamellipodia with a typical thickness of 100-150 nm, that is optimal for cryo-electron tomography (cryo-ET) analysis without altering the cellular integrity, or applying chemical fixation ^38^. This feature and the uninterrupted nature of the protrusive expansion on gal-8, enabled us to retrieve fundamental information on the assembly of the actin network in the expanding lamellipodium.

Here, we applied cryo-ET to mouse embryonic fibroblasts (MEFs) plated on gal-8-coated EM grids, enabling us to analyze the structure of the actin network at the cell expanding edges, *in cellulo*. We further used a combination of image processing approaches to determine quantitatively the polarity of individual actin filaments located at the protruding lamellipodium. These analyses revealed unexpected variations in the actin directionality that define functional sub-domains within the protruding lamellipodia of spreading cells, including a significant level of filaments that orient their barbed ends toward the cell center. These findings suggest that rapid reorganization of the actin network architecture occurs within the protruding leading edge of cells spread on galectins.

## Results

The 3D organization of the cells, including the interface between the actin cytoskeleton and the plasma membrane, was previously visualized by cryo-ET ^39^. However, a major limitation in this approach is the thickness of spreading cells, which hinders the spatial resolution and the precise localization of the cellular processes. These limitations can be overcome by allowing cells to spread over gal-8-coated surfaces which markedly reduces the thickness of the adhering cells ^37, 38^. It was demonstrated that the presence of gal-8 in the ECM correlates with cancer and metastasis ^40^, and increase cell growth and adhesion of metastatic cells ^41^.

### *In cellulo* structural analysis of actin and Arp2/3

MEFs cultured on gal-8-coated surfaces exhibited radial membrane protrusions within 20 min after contacting the coated surface (Fig. 1A and Supplementary Movie 1 and Movie 2). The spreading dynamics were significantly faster and continuous, compared to those obtained with other ECM proteins, e.g. fibronectin ^38^. However, the characteristic actin dynamics and treadmilling are detected (Supplementary Movie 2). Moreover, both Arp2/3 and F-actin showed a conspicuous association with the widely annular leading edge (Supplementary Fig. 1). In this study, we conducted experiments with cells growing on gal-8-coated EM grids that were vitrified within 10-20 min after engaged with the EM grids.

**Figure 1.**
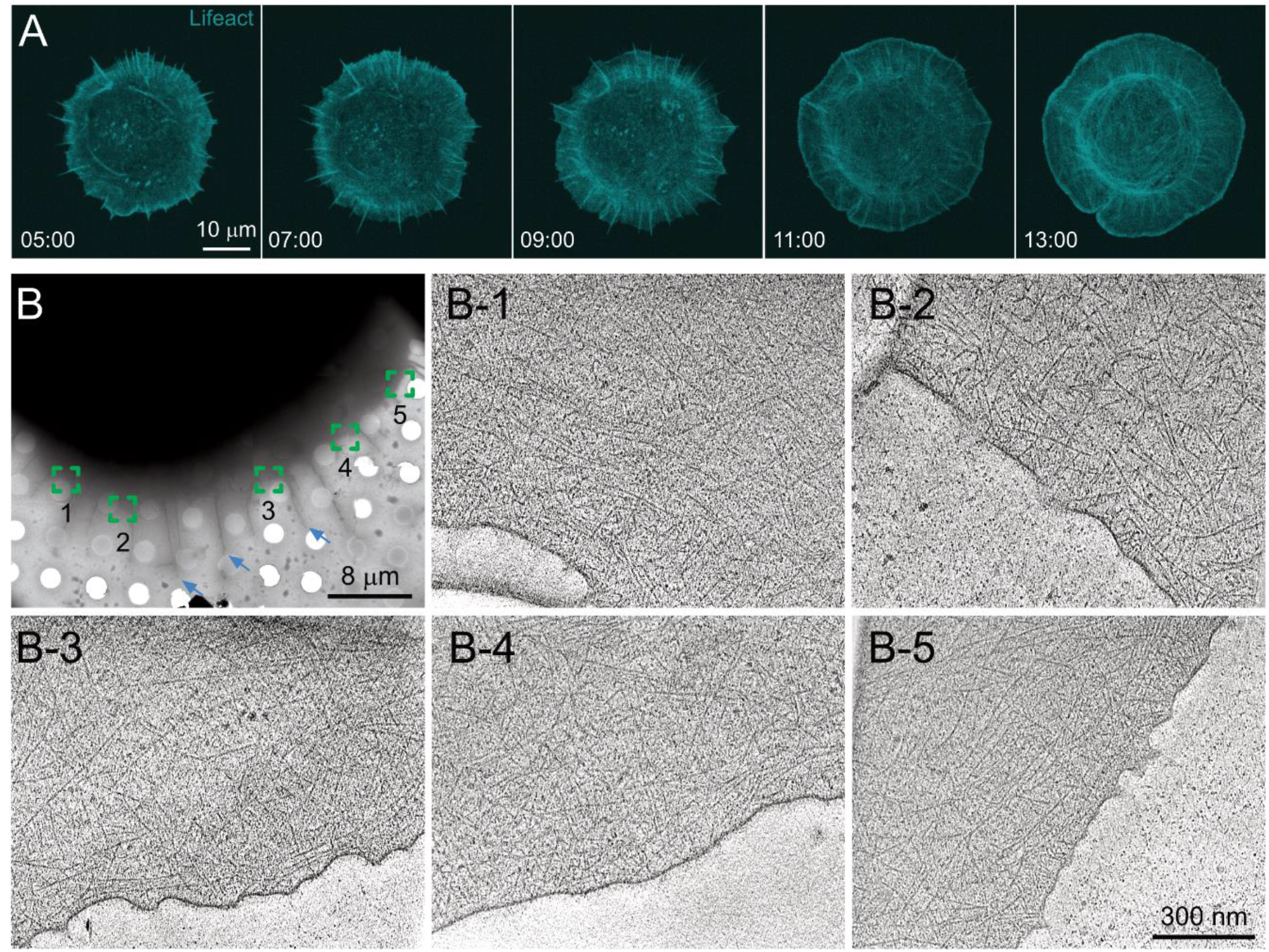
Cell spreading on galectin-8 coated substrate. **(A)** Time lapse images of a MEF cell transfected with Lifeact-mRuby let to spread on galectin-8 coated substrate (Supplementary Movie 2). **(B)** A cryo-EM image of a representative MEF cell spread on a galectin-8-coated EM grid. The area subjected to cryo-ET analyses indicated in boxes. A representative x-y slice through each tomogram in shown (**B-1** to **B-5**). Filopodia are indicated with cyan arrows.

Low magnification cryo-EM imaging revealed radially-spread cells, with both lamellipodial and filopodial protrusions (Fig. 1B). We primarily focused on the edges of the circumferential lamellipodia and acquired multiple tomograms. Fig. 1B-1 to 1B-5 show typical positions that were analyzed by cryo-ET. The thickness of the 57 acquired tomograms was 102 ± 25 nm, which provided us with an opportunity to systematically acquire high-quality data of essentially an entire peripheral ∼600 nm wide belt of the lamellipodium edge. Notably, when spread on fibronectin-coated grids, cells presented edges that are significantly thicker, typically ∼200 nm ^31, 42^.

Initially, actin filaments in 7 tomograms were manually segmented (Supplementary Fig. 2) and used for training a convolutional neural network ^43^, enabling us to automatically segment actin filaments from additional 50 tomograms. By applying the actin polarity toolbox (APT) ^44^, we extracted sub-tomograms and reconstructed the structure of the actin filament at 14.7 Å resolution (Fig. 2A-C and Supplementary Fig. 4A) determined by Fourier shell correlation to EMD-6179 ^45^. The 3D refined actin filament structure was used to determine the filament’s polarity throughout the analyzed volume.

**Figure 2.**
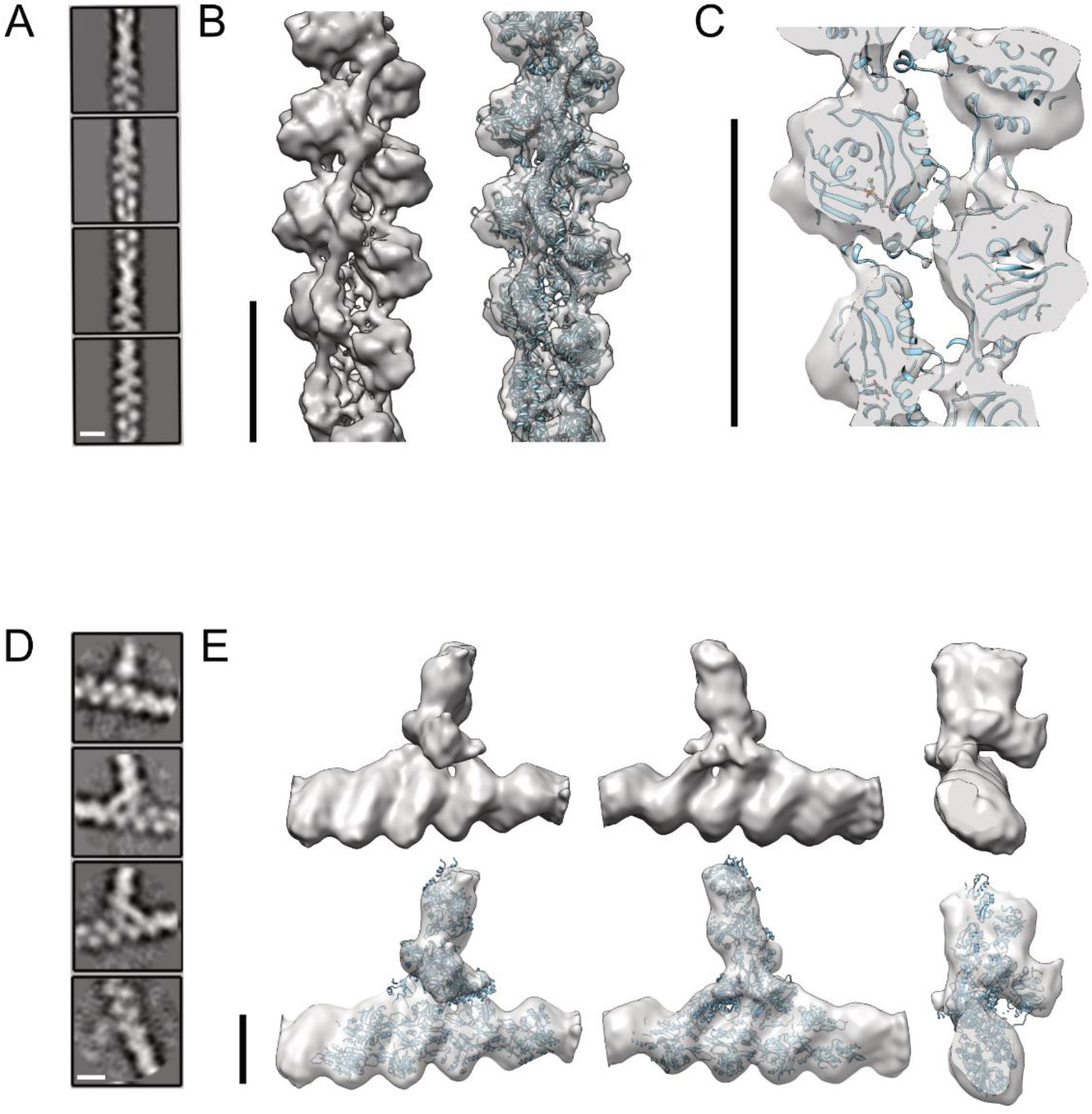
Three-dimensional reconstruction of actin and the Arp2/3 complex. **(A)** Representative 2D class averages of actin segments analyzed from cryo-tomograms of spreading cells. **(B)** The structure of actin filaments resolved to 14.7 Å (Supplementary Fig. 2A). Cryo-EM structure of actin EMD-6179 was docked into the structure calculated from in situ cryo-tomograms. **(C)** A cut away view through the *in situ* actin structure indicates the quality of the fit of the *in vitro* structure. **(D)** 2D structural classes of Arp2/3 mediated branches, as were identified by template matching (see methods section). **(E)** Reconstructed structure of actin branches at 26 Å resolution (Supplementary Fig. 2B). The structure of the extracted branches (EMD-11869) was docked into the *in situ* reconstructed structure (lower panel). Scale bar: 8 nm

The branched actin network of the lamellipodia is decorated by the Arp2/3 complex, which is found at branched junctions along the actin filaments ^46^. The Arp2/3 comprises the two actin-related proteins 2 and 3, Arp 2 and 3, in addition to ArpC 1,2,3,4, and 5 ^47^. This heptameric complex was reconstructed at 9 Å resolution using subtomogram averaging from cytoskeleton extracted cells ^48^, and *in-situ* to a more modest 32 Å resolution ^49^. Here, we utilized a template matching approach for localizing the complex in 96 tomograms (Supplementary Fig. 3A). We extracted sub-tomograms using a 31.8×31.8×31.8 nm^3^ box size, and applied 3D-subtomogram averaging. Fig. 2D shows class averages of projection images at different orientations in respect to the x-y plane, suggesting a reasonable orientation coverage. We obtained a refined 3D structure with a gold-standard resolution of 26 Å (Supplementary Fig. 4B) by using 6149 subtomograms. We were able to dock into our structure the Arp2/3 structure EMD-11869 (Fig. 2E), with a cross-correlation coefficient of 0.84 ^48^ analyzed using UCSF Chimera ^50^. The *in-situ* structural analysis of both actin filaments and the Arp2/3 allows to map these structures back into the reconstructed cellular volumes and investigate the organization of the lamellipodial actin network in high details.

### Mapping of actin filament’s orientation and polarity in the lamellipodia of spreading cells

The 3D network geometry of actin cytoskeleton affects the overall mechanical properties of cellular processes, using the asymmetrical polymerization dynamics of actin. The thermodynamic polarity of actin filaments, which relies on the existence of a fast-growing barbed end, plays a key role in regulating the dynamics and mechanics of the whole network. Developments in cryo-EM and tomography, in conjunction with novel image processing approaches, allow high-enough resolution to determine the precise polarity of individual actin filaments within the lamellipodial network. Initially, we visualized the complete organization of the actin filaments by isosurface rendering of the tomograms (Fig. 3A and Supplementary Fig. 2). This initial step provided the basis for comprehensive structural analysis of the orientation and polarity of the filaments within the tested volumes relative to the membrane. Towards this end, we mapped back the structure of the actin filaments to obtain the polarity information of each filament as reported (^44, 51^, methods section). We then quantified the relative angle between each actin filament and the closest plasma membrane by comparing the polarity of the filament to the normal angle of the nearest plasma membrane.

**Figure 3.**
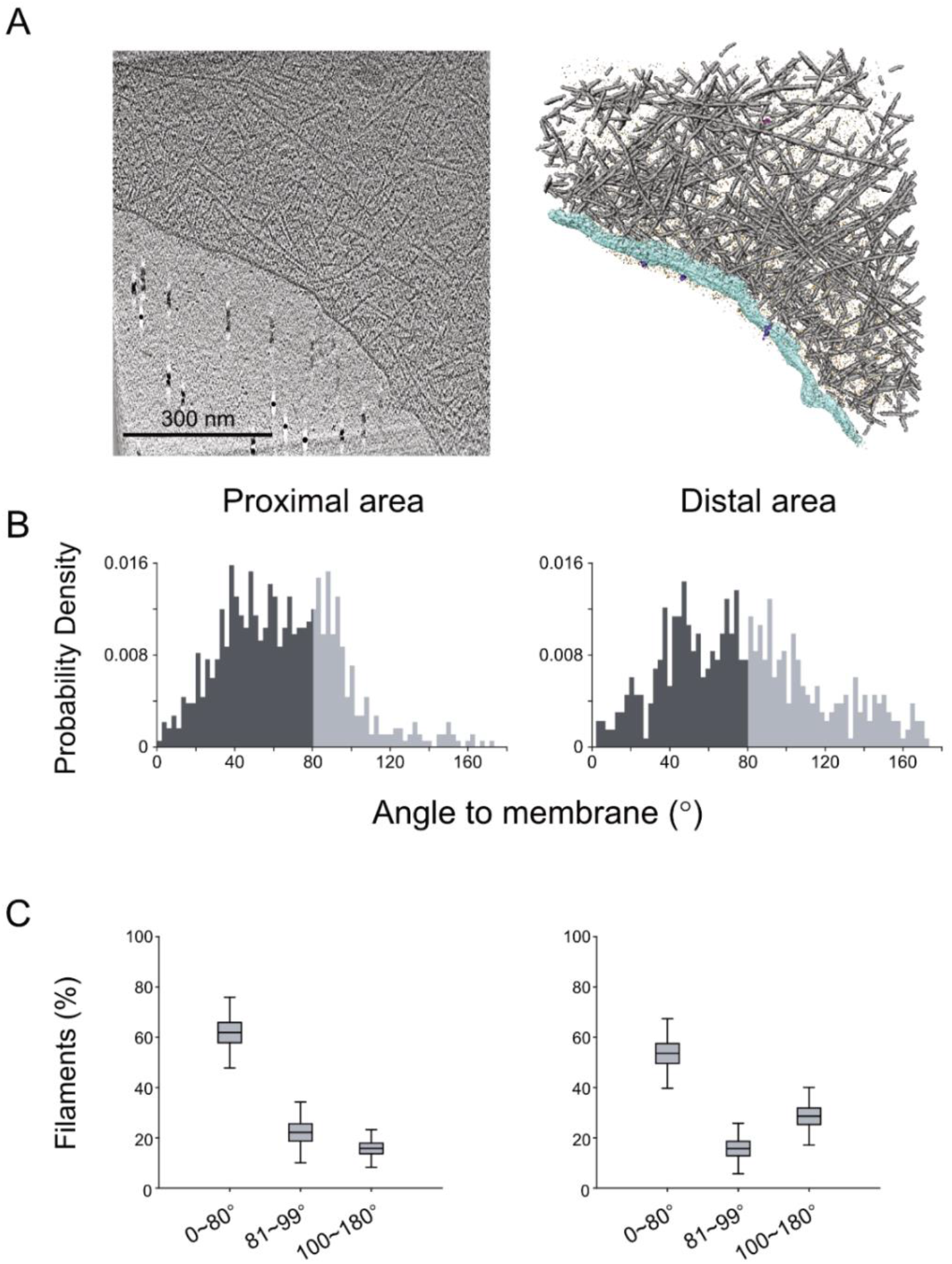
Filament network and filament-membrane orientation. **(A)** An x-y slice through a cryotomogram (left) and its rendered isosurface view of a lamellipodium protrusion. Actin filaments (gray), cell membrane (turquoise), membrane receptors (purple) pr and cell components (magenta). **(B)** The orientation of filaments in respect to the membrane was measured in 50 tomograms at the membrane proximal area of the cells (0-40 nm away from membrane) and distal area (360-400 nm away from membrane). The dark gray region represent filaments that contain vectorial components with the barbed end towards the membrane. **(C)** The boxplots showing the proportion of the different filament directionalities in respect to the membrane, with 1 SD and 1.96 SEM in whiskers. Color scheme is also used in Figure 4.

Initially, we compiled all the data and analyzed all filament-membrane orientations along the protruding membrane and with 40 nm slabs away from the membrane towards the cell center. The barbed end defines the representative coordinate of each filament (a polarity perpendicular and towards the plasma membrane is defined as 0°). In the membrane proximal area (0-40 nm away from the membrane), more than 60% of the filaments have their barbed ends towards the membrane (Fig 3B, left, dark gray). While in the membrane distal area (360-400 nm away from the membrane and towards the cell center), we found that 55% of the barbed ends are pointing towards the membrane (Fig. 3B, right, dark gray). Respectively, the density of the actin changes only moderately within the analyzed 400 nm distance from the membrane, with the highest filament density at ∼80 nm away from the membrane, (Supplementary Fig. 5A). The barbed end density shows a slight increase close to the membrane, presumably due to the higher nucleation activity at the membrane proximal area (Supplementary Fig. 5B). This shows that when merging data from a large number of lamellipodial volumes, neither the density nor the directionality of the actin seemed to change across the network, from the membrane towards the cell body. Thus, in such an analysis, radial and circumferential variations of actin density and polarity of potentially different network states may be averaged and hindered.

The actin cytoskeleton at the lamellipodia is branched by the Arp2/3 complex. In the acquired data, we found an Arp2/3 density of 2500 µm^-3^. Overall, the branched actin network showed a 1:3 ratio between the number of Arp2/3 and actin filaments (Supplementary Fig. 6A), suggesting that a significant number of actin filaments in the studied volumes were not branched.

### Radial diversity of filament orientations in the lamellipodia of spreading cells

To understand the organization of actin filaments and their orientation relative to the plasma membrane, we focused on the 400 nm-layer of the lamellipodial actin located underneath the plasma membrane of spreading cells, and analyzed the polarity of individual filaments as well as the 3D organization of the network. We found that the barbed ends of 60% of the filaments are oriented towards the membrane (forward orientation), 20% exhibit a parallel-filament orientation (defined as being largely parallel to the plasma membrane edge), while ∼10% point towards the cell interior (backward orientation) (Fig. 3C). We have defined the angular range of the three-characteristic orientations of actin filaments, relative to the protruding plasma membrane, as follows (Fig. 3C & 4A): ‘forward filaments’ are oriented at an angle of 0°-80° (with their barbed end towards the membrane) and shown in blue, the parallel filaments are oriented at an angle range of 81°-99° (mustard), while the ‘backward filaments’ are oriented at an angular range of 100°-180° (red). This color code allows for an easier appreciation of the overall network architecture while preserving the accurate orientation of individual filaments. We then compared the proportion of each of the 3 filament categories populating the first 400 nm-layer underneath the cell edge to identify the state of the lamellipodia.

**Figure 4.**
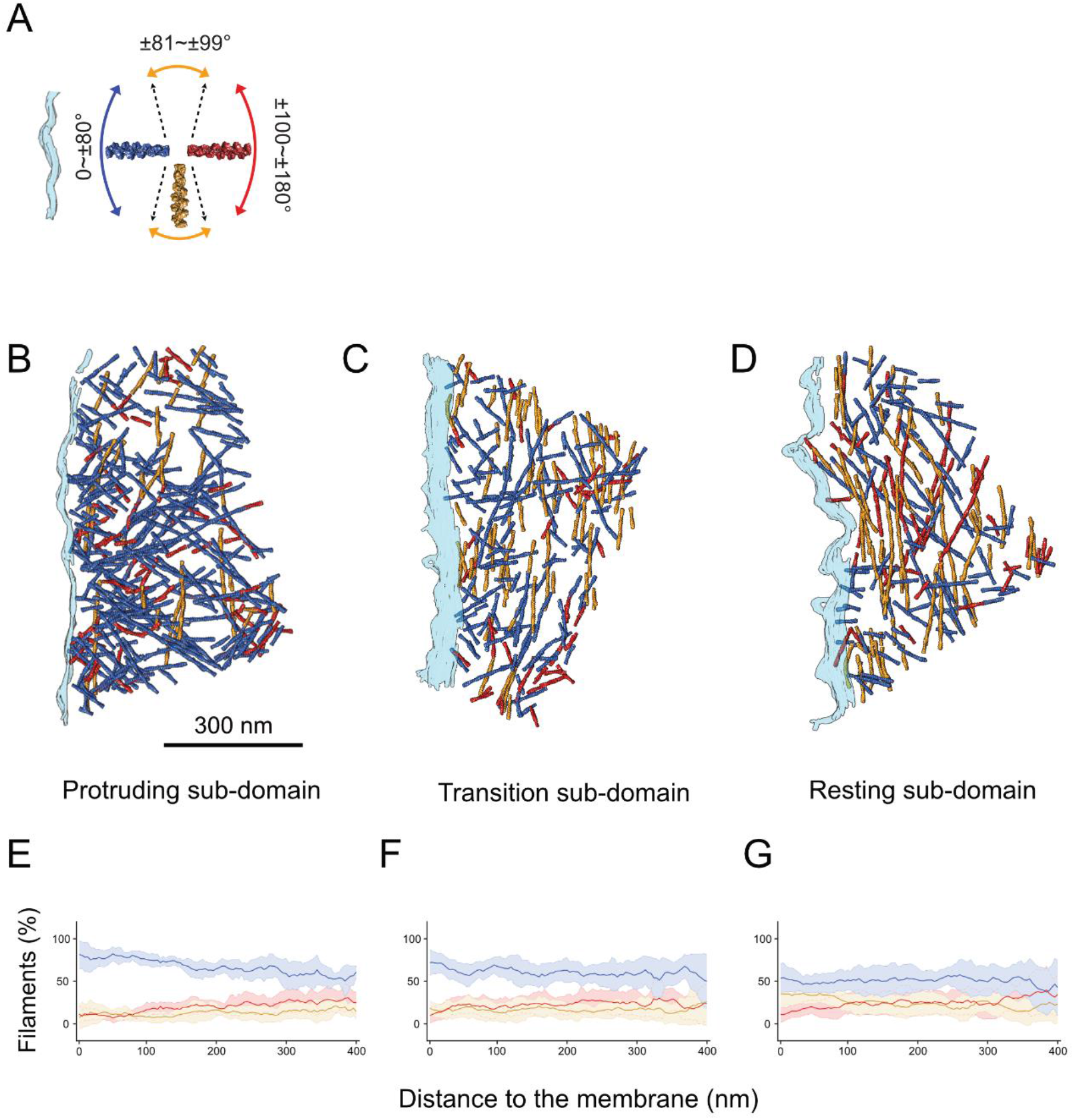
Three different sub-domains within lamellipodia. **(A)** A color scheme indicating the angles between actin filaments and plasma membrane (turquoise). Surface rendered views of representative tomograms from the 3 different sub-domains show the actin filaments colored according to their orientation in respect to the plasma membrane (**B-D)**. (**E-G)** The fraction of the forward (blue), parallel (mustard), and backward (red) actin orientations as a function of the distance from the membrane are plotted as continuous line with a shaded error bar, for protruding, transition, and resting sub-domains (N = 10, 16, 21).

Further confirmation of the assignment of actin polarity provided by comparison of the filament-to-membrane angle distribution of all the actin filaments to the distribution of Arp2/3, while each Arp2/3 is back-mapped from the averaged structure and assigned as a mother filament and a daughter filament. Both of Arp2/3 orientation and actin showed a similar angle distribution at around 80° (Supplementary Fig. 6C). By visualizing the branched network of Arp2/3 and actin filaments (Supplementary Fig. 3B), we confirmed that most actin filaments found in the Arp2/3 matches the known orientation of actin filaments.

### Circumferential diversity of filament orientations in the lamellipodia of spreading cells

Next, we have tested the circumferential variability of actin filament orientations along the expanding cell front. Our analysis revealed three main sub-domains within the ∼400 nm-wide belt of the spreading lamellipodium. Each sub-domain displayed a distinct pattern of actin organization. Based on the relative prominence of the forward actin filaments in these sub-domains we refer to them as the ‘protruding’, ‘transition’ and ‘resting’ sub-domains (Fig. 4B-D, Supplementary Fig. 7). The assignment of the sub-domain are neither biased by proximity to filopodia protrusions, nor can be predicted by the precise orientation of the plasma membrane (Supplementary Fig. 8). In the protruding sub-domain, the membrane proximal area exhibits the highest portion (around 85%) of forward-oriented filaments (Fig. 4E). The level reduced to about 60% in the most distal area (located at a distance of 360-400 nm away from the membrane) (Fig. 4E). Interestingly, the parallel-filaments display a comparable level in the proximal and distal areas of the protruding sub-domain, while the levels of backwards-filaments gradually increased from ∼10% in the proximal area to ∼30% in the distal area from the membrane (Fig. 4E).

In the ‘transition sub-domain’ (Fig. 4C), the network shows variable levels of forward filaments (in the range of 70% to 50%) that hardly correspond to their distance from the plasma membrane (Fig. 4F). Here, the prominence of the parallel filaments is higher than that detected in the protruding sub-domain, suggesting that while the network may still be protrusive, the parallel network and the backward filaments may support the establishment of contractile actomyosin force generation. This is more pronounced in the resting sub-domain, where around 40% of the actin filaments are parallel to the membrane (Fig. 4D and 4G).

A detailed comparison of actin directionality at the proximal and distal area (40 nm and 400 nm away from the membrane, respectively) is shown in Figure 5. Remarkably, in the protruding areas, the angle distribution of filaments at the proximal and distal areas varies significantly and broaden at the distal areas (Fig. 5A and 5B). In the proximal area, the highest prominent angle is ∼40°, while an almost even distribution ranging between 20-140° is found at distance of 400nm from the membrane. In the ‘transition sub-domains’, a bimodal distribution of actin directionality is found at the proximal area of the lamellipodia (Fig. 5C). A sub-population of the filaments is aiming at the membrane with an angle of ∼40° while a similar proportion of actin is oriented between 80°-90° (parallel orientation). Such an orientation has been suggested to support less protruding leading edge but would allow a dynamic growth of actin filaments that would support the emergence of a protruding network ^23, 52^. In contrast to the ‘protruding’ and ‘transition’ sub-domains, the proximal area of the ‘resting’ sub-domain (Fig. 5E) is characterized by highest frequent of parallel filaments. Surprisingly, throughout all sub-domains, we have found that a similar portion of backward oriented actin filaments with a slight increase in the distal area.

**Figure 5.**
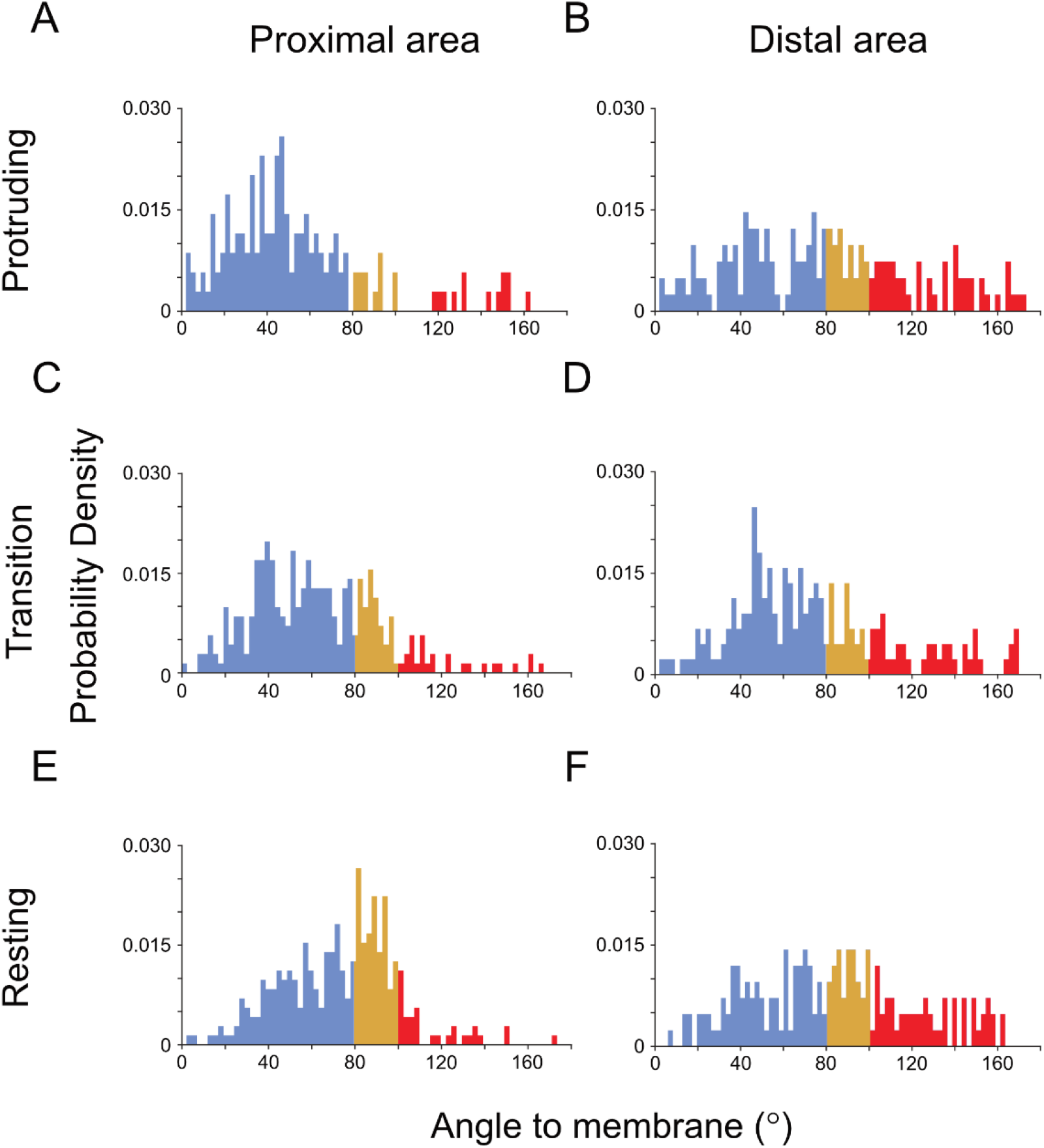
The distribution of filament orientations in the three different lamellipodia sub-domains. The orientation of actin filaments in respect to the plasma membrane is shown. The histograms show actin filament orientation at the membrane proximal regions and distal regions within the lamellipodia. Protruding (**A**,**B**), transition (**C**,**D**) and resting sub-domains (**E**,**F**). The filament-membrane orientation distribution at the membrane proximal (left) and distal areas (right). The colors indicate the angular orientation as shown in Figure 4A.

## Discussion

The reduction in lamellipodia thickness, induced by substrate-immobilized gal-8, allowed us to apply cryo-electron tomography for studying the molecular architecture of the actin cytoskeleton in the leading edge of intact spreading cells, at relatively high resolution. Gal-8 has shown to be a constituent of the ECM, functions as a pro-metastatic agent. It was suggested that it functions via enhanced cell adhesion properties ^53^.

Applying sub-tomogram averaging, we reconstructed the structure of the Arp2/3 complex, *in-situ*, and analyzed in detail the precise orientation and polarity of the actin network throughout the leading edge. This analysis revealed a heterogenous cytoskeletal organization, manifested by the presence of distinct structural sub-domains with different F-actin orientation, as summarized in Figure. 6. We propose that different actin organizations, both along the “radial axis” (proximal-distal, relative to the cell edge) and along the leading edge (“circumferential axis”) reflect aspects of the actin network dynamics. For example, there are regions in which the actin barbed ends are primarily directed towards the membrane (we refer to these regions as protruding sub-domains) while in neighboring regions there are prominence of filaments that run parallel to the plasma membrane, which we refer to as “resting sub-domains”.

**Figure 6.**
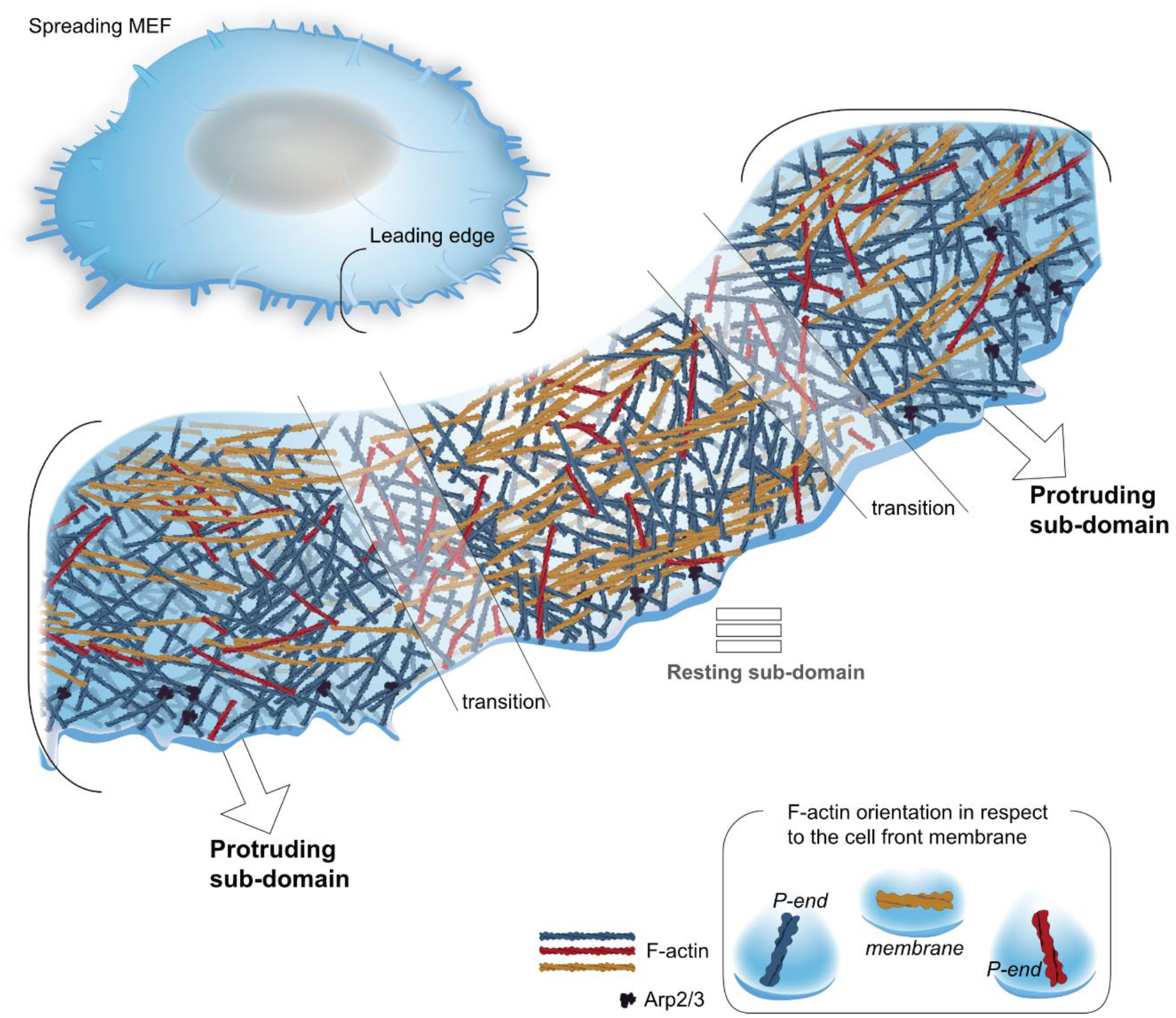
The lamellipodia architecture in spreading cells. A model depicting the actin organization at the edge of a spreading cell. Three distinct actin sub-domain exhibit different directionality of actin filaments were identified. A protruding sub-domain in which the majority of the actin filaments have their barbed ends pointing towards the membrane (blue), although parallel filaments (mustard) and filaments that are pointing towards the cell body (red) can be detected as well. In the resting sub-domain less filaments are directed towards the membrane while more parallel filaments are detected. Transition sub-domains are spaced between the protruding and resting domains. In all the three network architectures parallel oriented actin filaments (mustard) are enriched in the resting sub-domains although they can be seen in transition and protruding sub-domains as well. Similar amounts of filaments that are pointing towards the cell body (red) were detected in all sub-domains.

Additionally, the transition sub-domains, containing smaller fraction of parallel filaments than in the resting sub-domain, likely correspond to a transition state between protrusive and resting states. Surprisingly, in essentially all analyzed sub-domains, ∼10% of the filaments are oriented with their barbed end towards the cell center, rather than the expected opposite orientation, towards the plasma membrane ^24, 54-56^. The variations in filament organization were most pronounced along the radial axis, comparing regions that are located close to the protruding edge, to those located 300-400 nm away into the cell center. As indicated, the differences were primarily manifested by the relative prominence of forwards filaments oriented with their barbed ends towards the membrane along the radial axis ^23, 52, 54-56^ It is notable that the three sub-domains are in general agreement with previous reports which analyzed the actin network in leading edge of fish keratocytes and MEF cells ^23, 52, 57^. The data showed how changes in angles between actin filament and the plasma membrane can alter the mechanical load onto the network. Specifically, larger angles were shown to decrease the load and eventually lead to a reduced cell movement ^23, 52, 57^. Thus, filaments that encounter the membrane at larger angles might lag behind neighboring filaments that elongated centripetally toward the membrane. These stalled filaments may eventually detach from the membrane and produce a membrane parallel orientated filaments. Indeed, differentially oriented populations of actin meshwork were proposed to affect protruding and resting lamellipodia ^11^.

Our current observation, demonstrating a molecular heterogeneity in actin filament orientation and polarity, suggesting that within the lamellipodia of spreading cells there are sub-micron scale domains with distinct dynamic properties. This is apparently inconsistent with the view that actin polymerization in the leading edge is driving a seemingly uniform and coherent retrograde flow. It is unclear how individual filaments are changing their orientation while moving centripetally. Yet, it appears likely that molecular interactions of individual actin filaments with nearby immobile components, such as nascent focal complexes and diverse cytoskeletal complexes, introduce local perturbations to the retrograde flow, which create and reinforce the local heterogeneity at and near the leading edge. Such interactions of actin filaments may be fundamental for force generation that drive both the protrusion of the leading edge and the maturation of nascent integrin adhesions into mature focal adhesions ^58-60^. The typical average flow rate, as measured for example by direct fluorescence speckles microscopy ^61^, is in the order of 0.5-0.8 µm/min. However, the rate of this flow may vary spatially and temporally, due to local rates of polymerization, and the constant change in filament orientation. This may be too small and transient to be detected using live cell imaging. In fact, recent *in-vitro* and *in-situ* experiments demonstrated how actin meshwork, flowing over VBS1-activated vinculin can attenuate the flow, resulting in massive active bundling ^62, 63^. It is likely that actin filaments that encounter such local perturbations in live cells, undergo radical changes in their orientation while translocating from the ‘proximal zone’ (0-40 nm on the leading edge) to the ‘distal zone’, located ∼300-400 nm towards the cell center. As indicated within this timeframe, along the radial axis there is a decline in ‘forward filaments’ and an increase in the prominence of backward- and parallel-filaments.

A key component of the lamellipodium actin network, the Apr2/3 complex forms the branched actin filament arrays ^64^. Our findings, which are supported by previous observations, suggest rather low frequency of Arp2/3 branching as J.V. Small and colleagues measured the Arp2/3 branches in lamellipodia of fibroblasts ^26^. Using electron tomography of detergent treated and stained cells, they evaluated the branch density to one every 0.8 μm filament length. In this study, based on an unambiguous localization of Arp2/3 in intact cells, we detected genuine Arp2/3-mediated branches on only a fraction of the actin filaments within the lamellipodium, which translates to a branch every ∼0.6 μm of filament length. Since Arp2/3 mediated branch density affects the actin stiffness and is suggested to play a central role in lamellipodia protrusion, the density of branches may vary according to the cell type during faster or slower protruding processes ^65, 66^. Based on these findings we propose that structural fluctuations in protruding regions within the lamellipodia edge, may represent macroscopic manifestations of local variability in filament orientation and branching. This proposal is in agreement with previous studies suggesting limited (yet significant and reproducible) prominence of branches in protruding leading edges of fibroblasts ^26^.

In evaluating the significance of the results presented here to the cytoskeletal characteristic during cell spreading. With implications on cell migration, it is noteworthy that the spreading of cells on gal-8-coated substrates is continuous and overall faster than that on fibronectin which consists of repeated cycles of protrusions and retractions ^38^. Cell spreading and motility require the re-organization of the branched actin network into a mixed-polarity parallel bundles, a process that involves focal adhesion localized actin binders, such as vinculin and α-actinin ^62, 67^. The diverse actin orientations, which include actin directed with their barbed ends towards the cell center, create mixed polarity actin organization that allow an efficient establishment of contractile actomyosin complexes. Similarly, mixed polarity of actin was found in protruding platelets pseudopods ^51^.

## Martials and Methods

### Cell culture and sample preparation

MEFs expressing vinculin-venus ^44, 68, 69^ were used for optimal adhesion and spreading on EM grid. The cells were cultured in Dulbecco’s Modified Eagle’s Medium (Sigma-Aldrich, D5671) with 10 % fetal bovine serum (Sigma-Aldrich, G7524), 2 mM L-glutamine (Sigma-Aldrich, G7513) and 100 mg/ml penicillin-streptomycin (Sigma-Aldrich, P0781) at 37°C and 5% CO_2_. Glow-discharged EM grids with carbon support film (R2/2,Au mesh;Quantifoil, Jena, Germany) were coated with 25 mg/ml gal-8, 2hr room temperature. The cells were detached with 5mM EDTA, resuspended in serum-free medium, and seeded onto the EM grids. The cells were then incubated incubation at 37°C and 5% CO_2_ for 10 to 20 mins to allow cell spread. Before plunge-freezingin liquid ethane, the grids were washed with 1x PBS (Fisher Scientific, BP399-1), and 4 ml of BSA-coated 10nm fiducial gold markers (Aurion, Wageningen, Netherlands) was applied onto the grids.

### Immunofluorescence (IF) and live cell imaging

For live cell imaging, cells were plated at a density of 5 × 10^4^ cells ml^−1^ onto the 35 mm cell culture dish with 14 mm-diameter glass bottom (MatTek, catalogue number: P35G-1.5-14-C) coated with Gal-8. Video recordings started 2-5 mins after the cells were added to the dish. Interference reflection microscopy (IRM) time-lapse imaging were carried out using the DeltaVision RT microscopy system (Applied Precision Inc., Issaquah, WA, USA), equipped with a 100X oil immersion objective (1.3 NA, UPlanSApo), at 15 seconds time intervals between frames. Confocal images and videos were taken with ANDOR Dragonfly spinning disk confocal microscope using 100X objective and an sCMOS (Zyla) camera. The transfections were done with jetOPTIMUS^®^ (Polyplus-transfection^®^ SA), following the manufacturer protocols.

For IF, cells were plated on Gal-8 coated cover-slip and let spread for 16 min before fixation with 3.7% paraformaldehyde. After washed and permeabilized with 0.1% Triton-X/PBS for 3 times/ 5 mins, cells were blocked with 1% BSA in 0.1% Tween-20/PBS for 1 hr. Cells were then stained with 1:100 Anti-p34-Arc (Sigma-Aldrich, 07-227) for 1 hr. After 3 times wash of 5 mins with 0.1% Tween-20/PBS, cells were stained with 1:400 goat anti-rabbit IgG (Alexa Fluor® 488, Abcam) and 1:100 Alexa Fluor® 647 phalloidin for 1 hr. Before imaging, cover-slips were mounted with fluorescence mounting medium (Dako Omnis). Confocal images were taken with Olympus IXplore SpinSR10 super resolution imaging system equipped with two sCMOS cameras, and processed with ImageJ ^70^.

### Cryo-electron tomography

Titan Krios G2 transmission electron microscope (Thermo Fisher Scientific, Waltham, MA) equipped with energy filter and K2-summit direct election detector (Gatan, Pleasanton, CA) was used for data acquisition. The microscope was operated at 300 keV in zero-loss mode; energy filter slit width was set to 20 eV. The microscope was controlled by SerialEM ^71^. Tilt series were acquired ranging from - 60° to 60° with 3° increments at −4 µm defocus. The tilt series were acquired at dose-fractionated mode with a frame rate of 0.2 sec/frame to a total of 1.2 sec/projection. The magnification was 64,000x resulting in a pixel size of 2.207 Å with an accumulated electron dose of around 100 e^-^/Å^2^. The tomograms were reconstructed using the TOM toolbox ^72^ and CTF corrected ^73^. The data were collected from 5 different batches of experiments and at least 4 cells of each batch.

### Actin filaments polarity determination and 3D reconstruction

We followed the APT framework for filament polarity determination ^32^. In brief, 212,537 actin segments were extracted from 58 tomograms with an equidistant spacing of 11 nm along the filaments. The filaments were segmented manually with AMIRA (Thermo Fisher Scientific, Waltham, USA) and then automatically using a convolutional neural network ^43^. The automated segmentation was further processed in UCSF Chimera ^50^ using the hide dust command and trimming the volumes to reduce false positive labeling. The subtomograms box size was set to 36 nm^3^. Next, the central 11 nm of each subtomogram were projected prior to 2D classification (Fig. 2A) and 3D reconstruction by RELION ^74^. For the final reconstruction of F-actin, 149,617 particles were selected and 3D refined, prior to the polarity determination.

To determine the polarity of the actin filament network, we adapted the APT for dendritic actin networks and the determination of filament-membrane orientation. In each tomogram, actin filaments were defined as 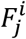 with *i* the filament number and *j*. the segment number within the filament. Actin filaments are generally capped and branched at the leading edge. To precisely define each filament at the branches, connected filaments must be segmented separately at the branching joints. Here, we fragmented each filament *i* by the correlation of the 3D coordinates of each segment *j*. If the next coordinate *N* + *1* cannot follow the tendency of the previous two coordinates 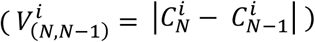 by 90 degree, the filament will be truncated and an additional filament will be generated from the break point deg 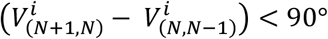.

Subsequently, we compared the polarity of each segment 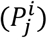 to the overall polarity of the filament (*S*(*F*^*i*^)). As actin filaments are in general short and straight within the tomogram, we define the overall polarity of the filament with the coordinates of the first and the last segment 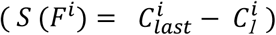. The segments following the overall polarity within 30 degrees were given a direction labeled *1* and else *0* 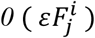;

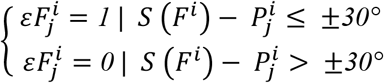

The *mean*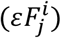 were used for the measurement of the filament polarity and the confidence of the polarity assignment was assessed based on 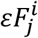 (see further detail in ^44^. Finally, only filaments passed the confidence test (combined confidence score (*ccs*(*F*^*i*^) > *0.6*), Supplementary Fig. 4C) were then used for actin network analysis.

### Filament-membrane orientation

The cell membrane of each tomogram *t* was manually segmented into *j* segments 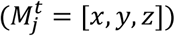 and defined the local normal vectors 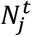 by a plane-fit to the adjacent membrane points within a sphere with a radius of 40 nm. The normal vectors were then smoothed with the adjacent normal vectors 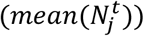. We then fetched the closest membrane points for each actin segment. The membrane-segment orientation of each actin segment 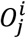 was then calculated by comparing the segment polarity and the average membrane normal vector 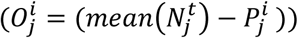. Finally, we averaged the membrane-segment orientation within a single filament to calculate its membrane-filament orientation 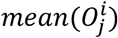.

For general categorization of the membrane-filament orientation, we defined the membrane-filament orientation into 3 intervals:

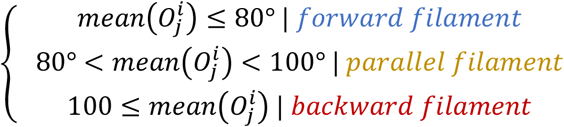

### Quantification and Statistical analysis

For categorization of different stages of lamellipodia, the proportion of forward, arc and backward filaments along the distance to the membrane were used (Fig. 4E-4G). In every tomogram, the proportion of the 3 filament types was continuously calculated with 100 frames (frame size 40 nm). To reduce outliers effect, we used robust fit function in MATLAB for estimation of the proportion of the 3 filament types in every tomogram. The estimated filament proportion at the first frames in all the tomograms were then used for identifying different lamellipodia stages. The portion of forward, transition, and backward filaments at the first frame were used as the coordinates of each tomogram for *k-means* clustering ^75^. We clustered the tomograms into 3 groups. The combined filament proportions were directly used for continuous linear plotting (Fig. 4E-4G). Histogram were normalized with the probability density function 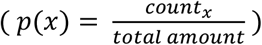 (Fig. 3B and Fig. 5). Continuous linear plots with error bars were plotted with the mseb function ^76^ (Fig. 4E-4G and Supplementary Fig. 5) and box plots (Fig. 3C and Supplementary Fig. 6) with the notBoxPlot function ^77^ in MATLAB.

### Tomogram visualization

All isosurface visualizations of actin filaments were rendered using UCSF Chimera or AMIRA. For visualization of actin filaments in Fig. 4B-4D and Supplementary Fig. 3B and Supplementary Fig. 7, the refined 3D F-actin structure was used for representing. The refined 3D Arp2/3 structure was used in Supplementary Fig. 3B.

### Template matching and 3D averaging of Arp2/3

We used the Arp2/3-actin structure as a reference (EMD-4790, ^49^) with a reported resolution at 32 Å for template matching procedure as described before ^78^. For that purpose we used 57 tomograms. The subtomograms were extracted with a box size of 31.8 nm. We then projected the subtomograms into 2D and performed 2D classification in Relion ^74^. 1,004 particles were selected and 3D averaged for the *de-novo* 3D structure. We then expanded the data to 95 tomograms and performed template matching again with the new reference. After subtomogram projection and 2D classification, the selected 2D images were aligned to a template library generated by rotating the reference in 6° increment. We then performed 2D classification with restricted searching angle to 2° and mask around the mother filament and branch. The selected 2D images were then back-mapped to the tomograms and manually cleaned up by the reasonable particle coordinates in IMOD ^79^. With a final 2D classification after the manual clean-up, 6,149 particles were selected for 3D averaging in PyTom^78^.

## Acknowledgements

We thank the center for microscopy and image analysis (ZMB) at the University of Zurich.

## Author contributions

W.L.C. prepared the acquired the tomograms,light microscopy images, and analyzed the data with the help of M.E. R.B.P. contributed to cell culture and protein purifications. W.L conducted the cell spreading assays and movies. B.G and O.M. conceived the work and financed the project. O.M wrote the manuscript with contributions from all authors.

