## Supplemental figures for "A network of mixed actin polarity in the leading edge of spreading cells"

### Supplementary Material

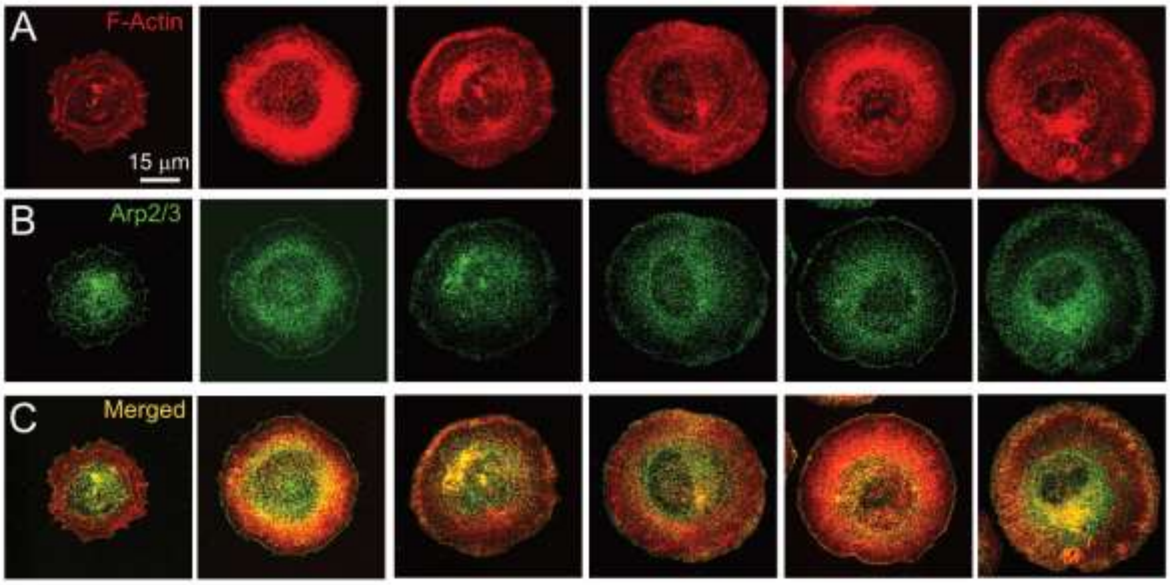


**Supplementary Fig. 1**. **Lamellipodia in cells, spreading on Galectin-8.** Immunofluorescent microscopy of MEFs spreading on Gal-8 coated substrate were acquired with spinning disk confocal microscopy. The Z-stack images were projected with ImageJ. The cells were chemically fixed 15 min after engaging to the Gal-8 coating glass. The six cells that are shown were stained with **(A)** phalloidin and **(B)** anti-p34-Arc. Merged color images are shown in **(C).**

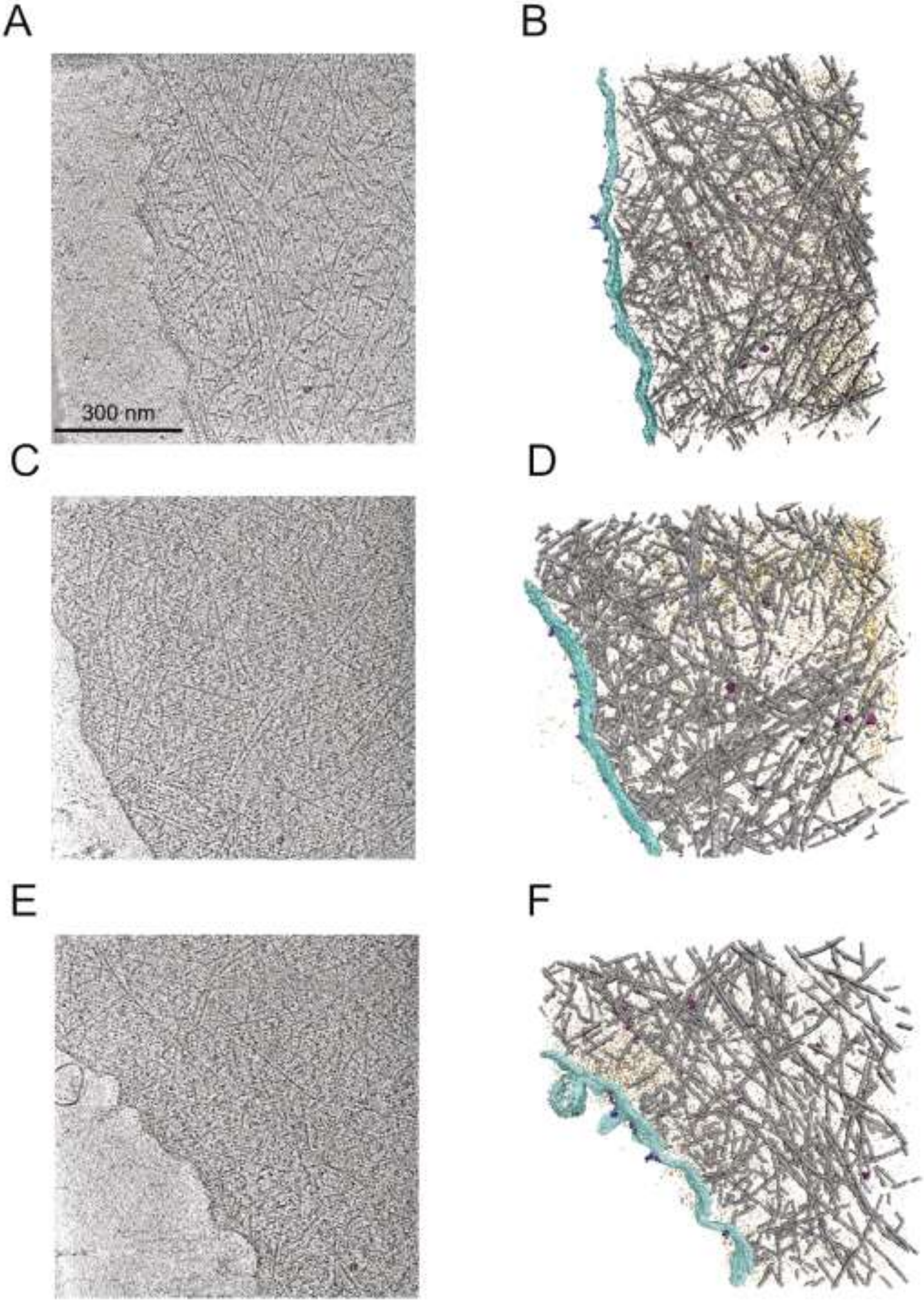


**Supplementary Fig. 2. Cryo-tomograms of lamellipodia of MEFs spread on Gal-8.** Cryo-tomograms of three cells are shown. x-y slices, 35.6 nm in thickness, through the tomogrmas (A,C,E) and the respective rendering isosurface views of the cryo-tomograms (B, D, F). Acin filaments (gray), membrane (turquoise), receptors (purple) and macromolecular complexes (dark red) are shown.

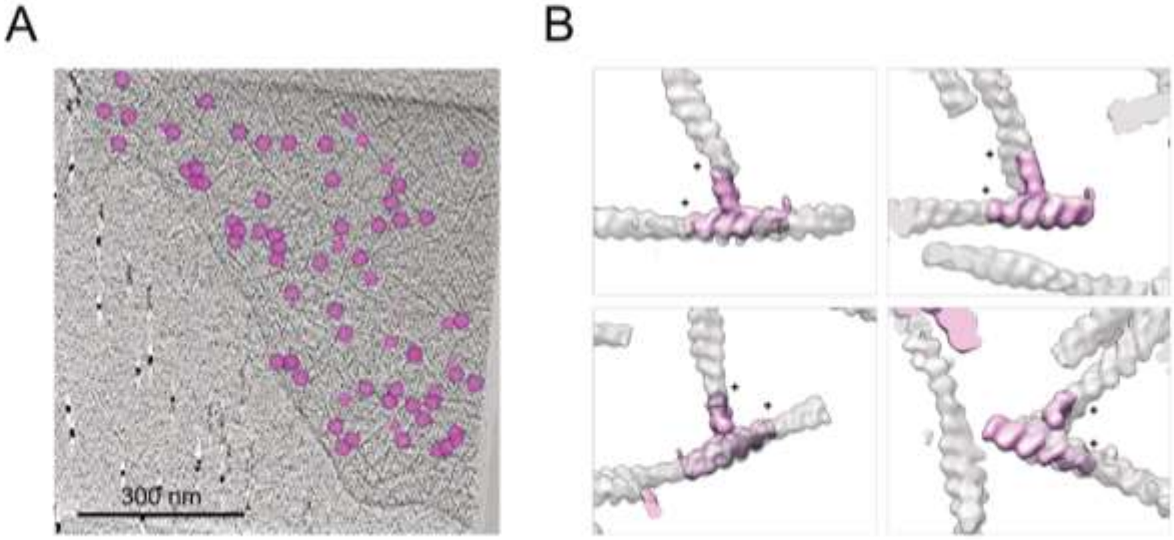


**Supplementary Fig. 3. The localization of Arp2/3 identified by template matching. (A)** The coordinates of the Arp2/3 (magenta) were superpositions on a 8.9 nm thick, x-y slice through the tomogram. **(B)** Surface rendered views of actin branches. Arp2/3 complexes are in magenta and actin filaments are in translucent gray. Barbed ends of daughter and mother filaments are marked with plus symbols.

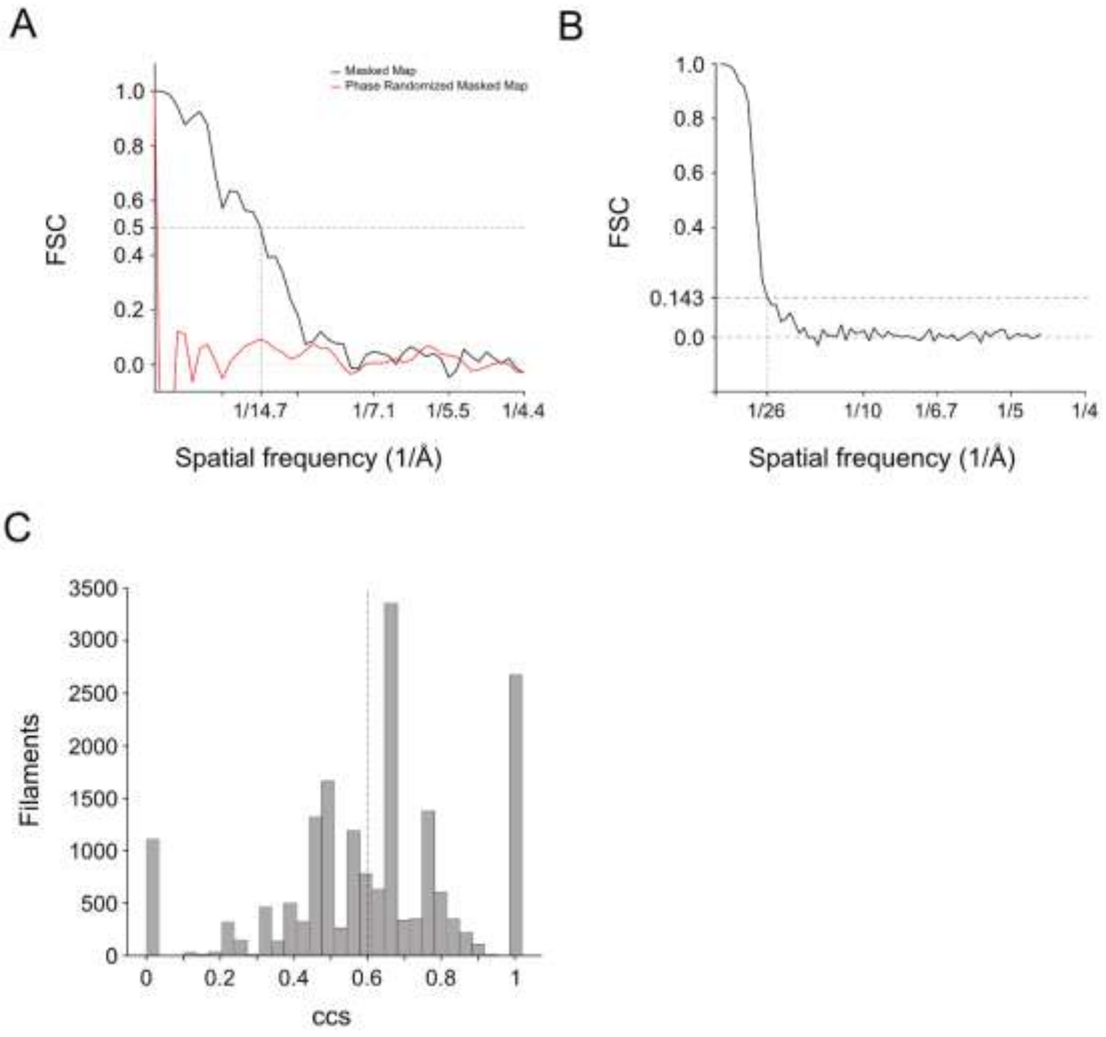


**Supplementary Fig. 4. (A)** The refined actin structure in Fig. 2B and 2C shows spatial frequency of 1/14.7 Å indicated by 0.5 of Fourier shell correlation to EMD-6179. **(B)** The refined Arp2/3 structure in Fig. 2E shows spatial frequency of 1/26 Å indicated by 0.143 gold-standard Fourier shell correlation criteria. **(C)** Combined confidence score (ccs) of the acquired data described in method. ~70% of the filaments have passed the 0.6 ccs threshold.

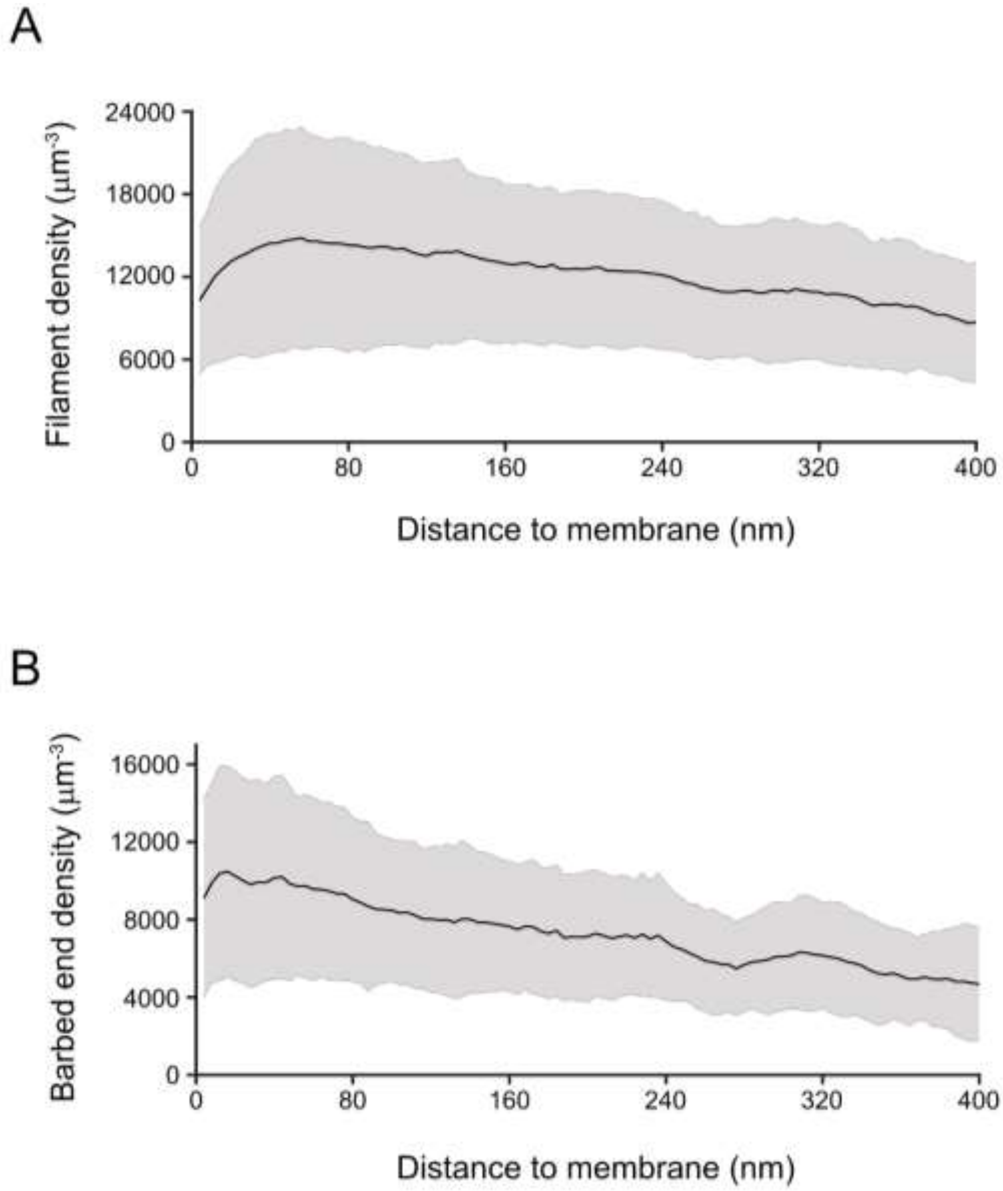


**Supplementary Fig. 5. (A)** Continuous filament density with shaded error bar along the distance to the membrane (N = 38). **(B)** Continuous barbed end density with shaded error bar along the distance to the membrane (N = 39).

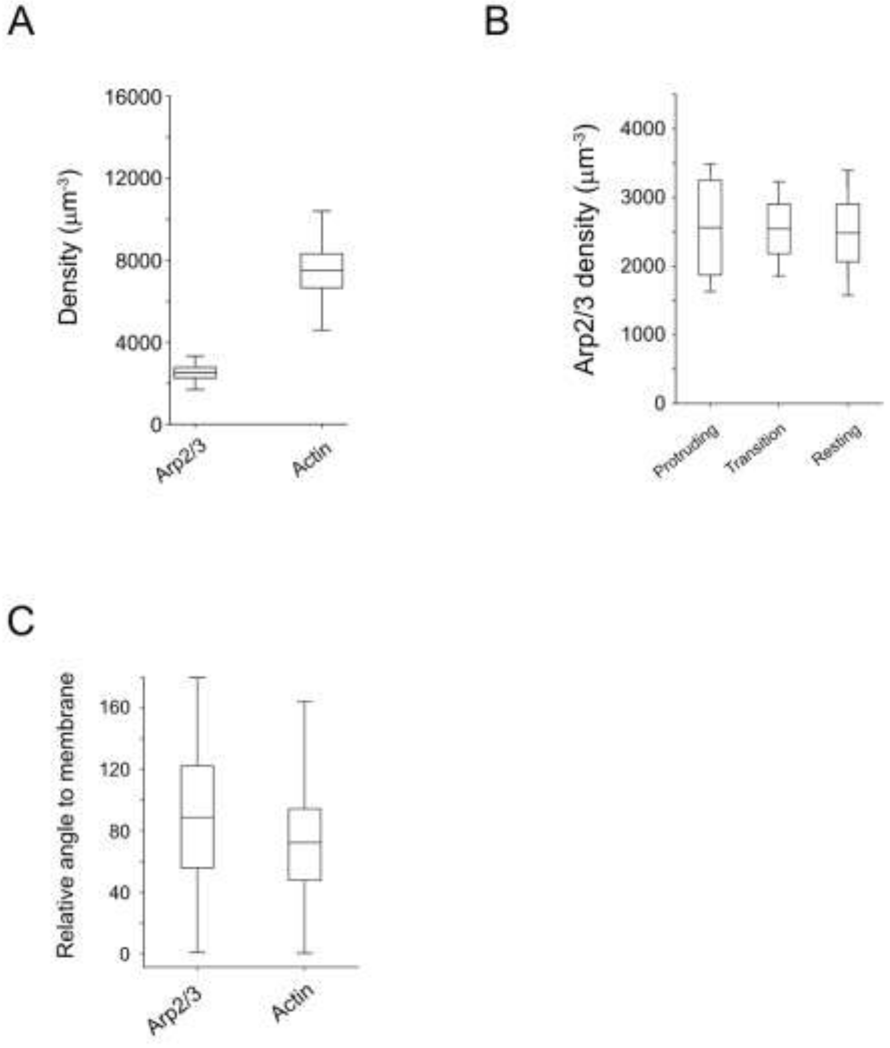


**Supplementary Fig. 6. (A)** Boxplot shows the density of actin and Arp2/3 with 1 SD and 1.96 SEM in whiskers. **(B)** Boxplot shows the Arp2/3 density in three sub-domains with 1 SD and 1.96 SEM in whiskers. **(C)** Boxplot shows the median and interquartile range of relative angle to membrane of actin filament and Arp2/3. Whiskers show the maximum and minimum. (N = 40)

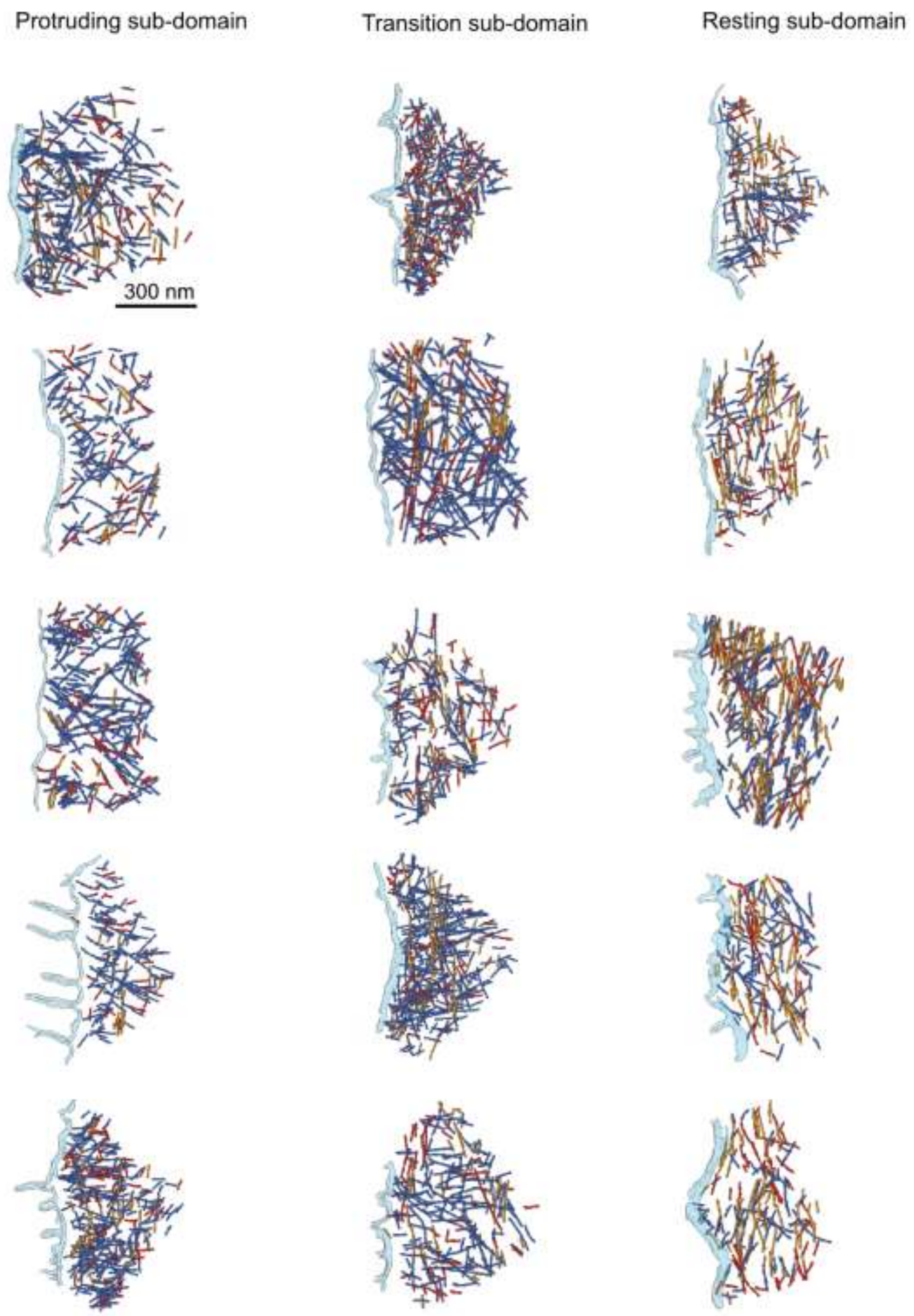


**Supplementary Fig. 7.** A collection of rendered tomograms, representing the 3 lamellipodial sub-domains.

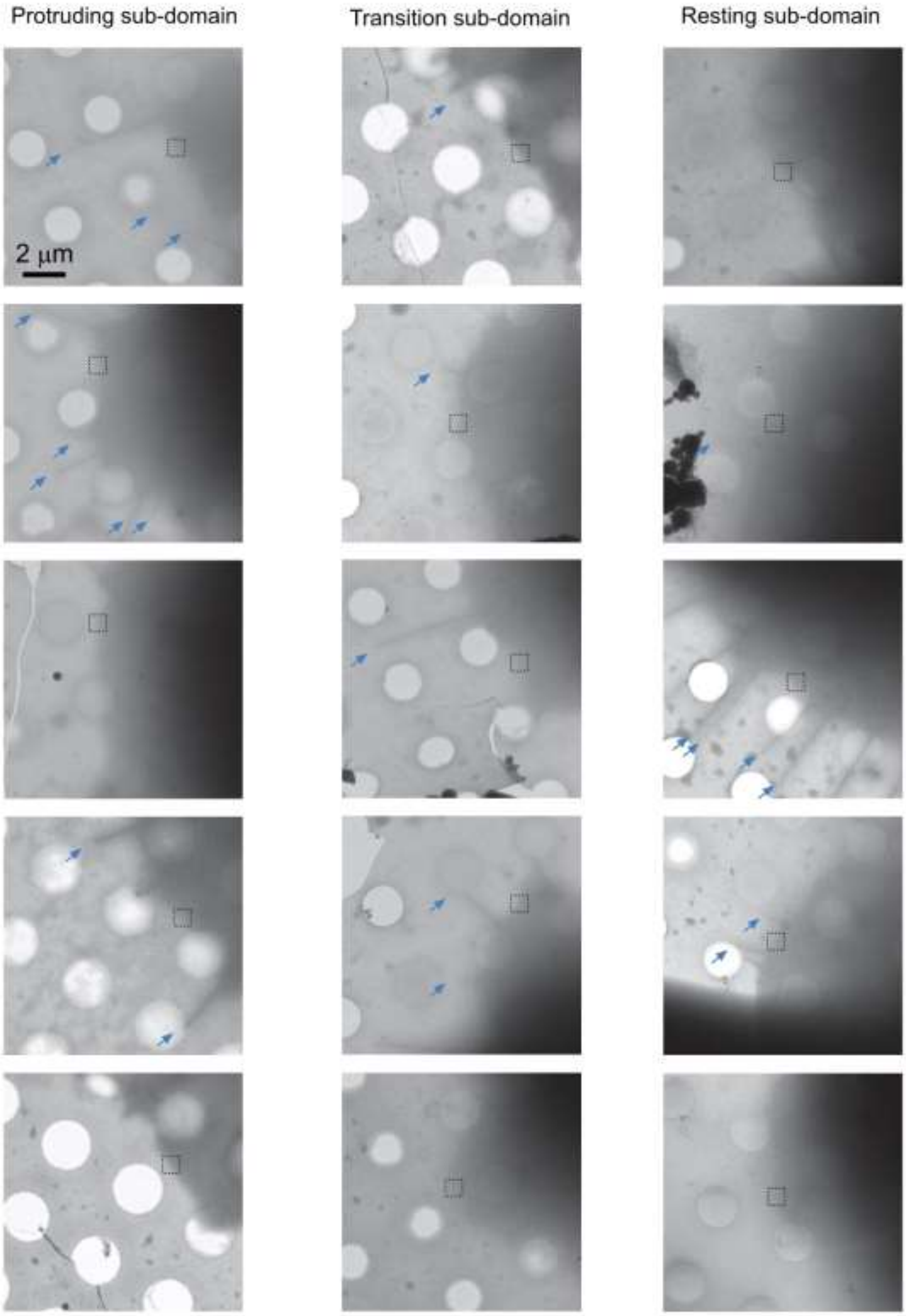


**Supplementary Fig. 8.** A collection of low magnification images of cell border. The box areas correlate to the data collected area in Supplementary Fig. 7. Filopodia are indicated with cyan arrows.

**Supplementary Movie 1.** A representatively time-lapse series of MEF cell spreading on galectin-8 coated substrate, imaged by interference reflection microscopy.

**Supplementary Movie 2.** A representatively time-lapse series of MEF cell spreading on galectin-8 coated substrate. The cell was transfected with Lifeact-mRuby.
